# SHIMS 3.0: Highly efficient single-haplotype iterative mapping and sequencing using ultra-long nanopore reads

**DOI:** 10.1101/2020.09.18.303735

**Authors:** Daniel W. Bellott, Ting-Jan Cho, Emily K. Jackson, Helen Skaletsky, Jennifer F. Hughes, David C. Page

## Abstract

The reference sequence of structurally complex regions can only be obtained through a highly accurate clone-based approach that we call Single-Haplotype Iterative Mapping and Sequencing (SHIMS). In recent years, improvements to SHIMS have reduced the cost and time required by two orders of magnitude, but internally repetitive clones still require extensive manual effort to transform draft assemblies into reference-quality finished sequences. Here we describe SHIMS 3.0, an extension of our SHIMS sequencing strategy, using ultra-long nanopore reads to augment the Illumina data from SHIMS 2.0 assemblies and resolve internally repetitive structures. This greatly minimizes the need for manual finishing of Illumina-based draft assemblies, allowing a small team with no prior finishing experience to sequence challenging targets with high accuracy. This protocol proceeds from clone-picking to finished assemblies in 2 weeks for about 80 dollars per clone. We recently used this protocol to produce reference sequence of structurally complex palindromes on chimpanzee and rhesus macaque X chromosomes; as a further demonstration of the capabilities of SHIMS 3.0, we finish the TSPY array on the human Y chromosome, which could not be resolved by previous sequencing methods. Our protocol provides access to structurally complex regions that would otherwise be inaccessible from whole-genome shotgun data or require an impractical amount of manual effort to generate an accurate assembly.

## Introduction

### Background and applications

Reference genome sequence quality is of central importance to modern biological research. Experiments based on aligning cheap and abundant short reads to existing reference sequences have become commonplace, permitting studies of variation by genome and exome resequencing, transcription by RNA sequencing, and epigenetic modifications by chromatin immunoprecipitation–sequencing. However, these experiments are limited by the quality and completeness of the underlying reference sequence, so that new insights may emerge from reanalyzing short-read datasets in the light of an improved reference sequence. The foremost obstacles to accurate reference genome assembly are repeated sequences within the genome. The most structurally complex repeats are ampliconic sequences – euchromatic repeats with greater than 99% identity over more than 10 kb^1^. The complex repetitive structures in amplicons mediate deletions, duplications, and inversions associated with human disease^2^. Amplicons pose special challenges for genome assembly, requiring extremely long and accurate reads to discriminate between amplicon copies and produce a correct reference sequence.

We developed our Single Haplotype Iterative Mapping and Sequencing (SHIMS) approach to cope with the ampliconic sequences of the human Y chromosome^3^. Because paralogous ampliconic repeats are more similar than alleles, we sequenced large-insert clones from a single haplotype, allowing us to confidently identify the rare sequence family variants (SFVs) that distinguish paralogous repeats in highly accurate (less than 1 error per megabase) synthetic long reads^3^. Mapping and sequencing were coupled; newly sequenced clones provide novel SFVs that refine the clone map and serve as markers to select new clones. SHIMS has been instrumental to producing reference sequences of structurally complex sex chromosomes from several species^4–10^, as well as the human immunoglobulin gene cluster^11^, and other structurally complex regions on human autosomes^12^. SHIMS remains the only sequencing approach that can reliably disentangle ampliconic repeats. Whole genome shotgun (WGS) strategies are constrained by a tradeoff between read length and accuracy among existing sequencing technologies. Sanger or Illumina reads are accurate, but are not long enough to traverse interspersed repeats, much less ampliconic sequence^13^. Single-molecule sequencing technologies like PacBio or nanopore sequencing offer reads long enough to span interspersed repeats and smaller ampliconic sequences, but lack the accuracy to disentangle nearly identical ampliconic repeats^14^. As originally implemented, SHIMS 1.0 required the resources of a fully-staffed genome center to generate Sanger reads, assemble draft sequences, and manually ‘finish’ each clone. We developed SHIMS 2.0 to combine the advantages of a hierarchical clone-based strategy with high-throughput sequencing technologies, allowing a small team to generate sequence, while reducing time and cost by two orders of magnitude, while maintaining high accuracy^15^. However, SHIMS 2.0 still required intensive manual review to resolve internally repetitive clones, and in some cases – particularly short, nearly perfect, tandem repeats – complete resolution remained impossible.

Here we describe SHIMS 3.0, an extension of our SHIMS sequencing strategy that we recently employed to produce reference sequence of structurally complex regions of chimpanzee and rhesus macaque X chromosomes^10^. SHIMS 3.0 uses a combination of nanopore and Illumina sequencing technologies to resolve repetitive structures within individual large-insert clones. We describe a protocol for generating full-length nanopore reads for pools of clones, and combining the structural information from these full-length reads with highly accurate short-read data to automatically produce assemblies of internally repetitive clones (Fig. 1). This protocol proceeds from clone-picking to finished assemblies in 2 weeks for about 80 dollars per clone, an improvement of 2 orders of magnitude compared with 24 months and 4000 dollars under SHIMS 1.0. As further demonstration of the performance of SHIMS 3.0, we apply this protocol to resolve the TSPY array on the human Y chromosome. The TSPY array is one of the largest and most homogeneous protein-coding tandem arrays in the human genome^16^, and it could not be completely resolved in the SHIMS 1.0 reference sequence of the human Y chromosome^4^.

**Figure 1.**
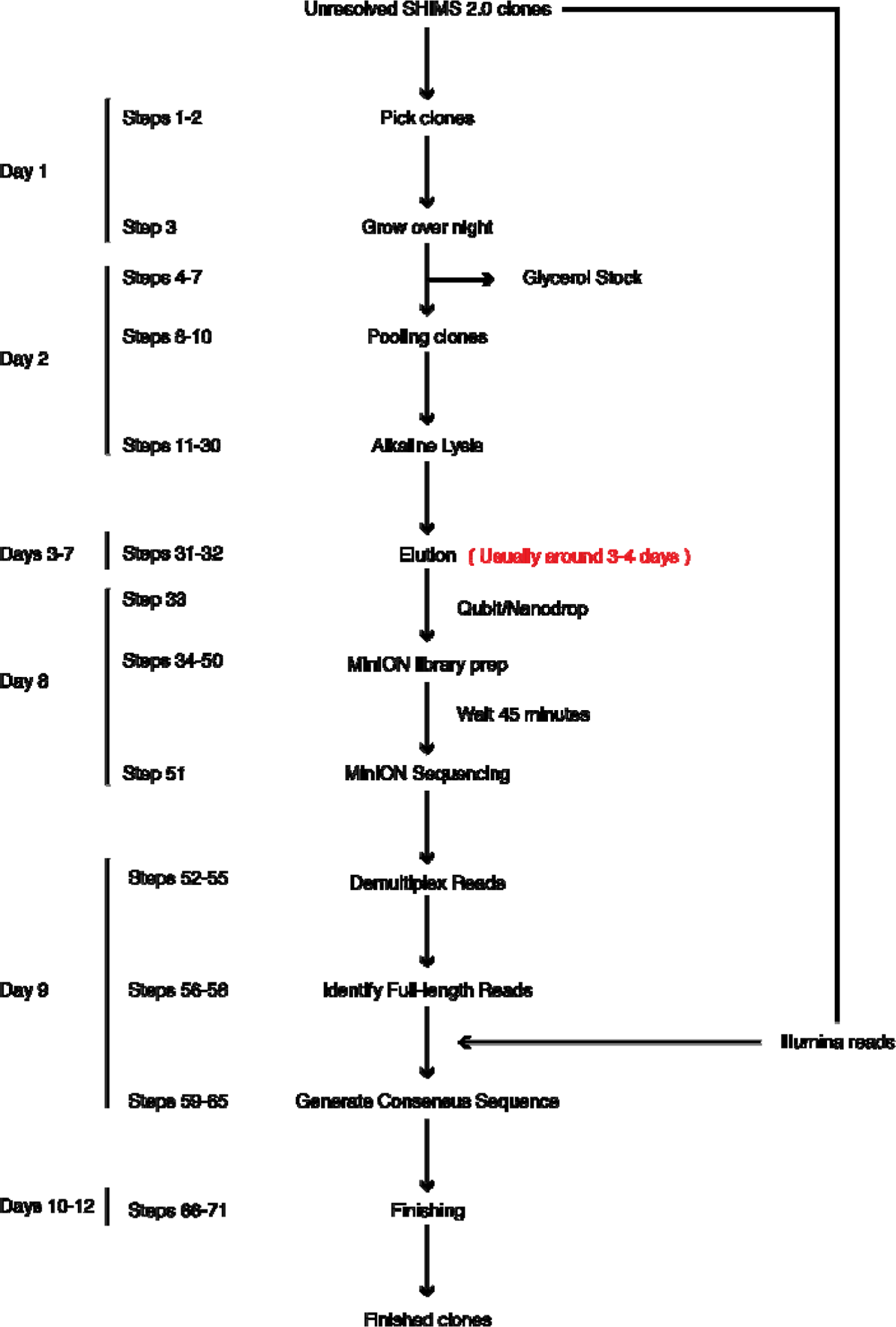
Overview of the SHIMS3.0 protocol. A timeline of a single iteration of the SHIMS 3.0 protocol, showing the major protocol steps, with key quality controls on the right. During a two-week iteration, 24 clones are processed in parallel to rapidly generate finished sequence from structurally complex clones. A single technician can proceed from a list of clones to full-length nanopore libraries in 8 d. After a brief MinION run overnight, a bioinformatics specialist can demultiplex fastq sequences, identify full-length reads, then polish and edit the consensus of these reads to generate finished clone sequence.

### Methodology

Large-insert clone libraries derived from a single haplotype are essential to the SHIMS strategy, and are discussed in detail in our description of SHIMS 2.0^15^. In brief, any library derived from an individual of the heterogametic sex will provide a single haplotype source for sequencing sex chromosomes, albeit at half the coverage of the autosomes. Libraries created from inbred strains can provide a single-haplotype source for autosomes. When inbreeding is not possible, special measures may be necessary to obtain a single-haplotype source of DNA^12^. The ideal library will have greater than 10x coverage of the chromosome of interest to minimize the number of gaps in library coverage. In SHIMS 1.0 and 2.0 it was important to match the average library insert size to the expected amplicon unit size, such that it was rare for two units to be present in the same clone^15^. SHIMS 3.0 uses ultra-long nanopore reads to span entire BAC clones^17^; therefore, we now recommend striving for the largest possible insert size, to minimize library construction, screening, and sequencing costs, as fewer clones will be necessary to achieve the required level of coverage.

All SHIMS strategies begin by selecting an initial tiling path of large-insert clones for sequencing and iterative refinement. Depending on the resources available for each library, it may be possible to identify clones of interest electronically, using fingerprint maps or end sequences; screening high-density filters by hybridization with labeled oligos, or high-dimensional pools for sequence-tagged-site content by PCR. It is most cost-effective to confirm the identity of each clone by generating draft sequence with the SHIMS 2.0 protocol, rather than designing specific assays for each clone^15^. In brief, this highly parallel method involves shearing BAC DNA to generate large (∼1 kb) fragments for individually indexed Illumina TruSeq– compatible libraries to sequence and assemble pools of 192 clones in a single week.^15^ In our experience, the structure of ampliconic regions is often unclear until a nearly complete tiling path is assembled, as the sequence map gradually unfolds as new variants are identified by sequencing. It is therefore preferable to seed the first iteration with as many clones as possible to identify sequence family variants early, and minimize the total number of iterations.

Draft clone assemblies generated from Sanger or Illumina reads are accurate enough to identify sequence family variants, and identify a minimum tiling path of clones. In previous iterations of SHIMS, each clone in this path would be painstakingly ‘finished’ to produce as correct and contiguous a sequence as possible. Highly skilled technicians would inspect draft assemblies for errors and anomalies, order and orient all draft sequence contigs, close all gaps, and resolve or annotate all sequence ambiguities (e.g. SSRs). SHIMS 3.0 departs from this approach, instead relying on the ability of nanopore-based sequencing technology to generate full-length reads to scaffold short-read assemblies and eliminate the need for laborious and time-consuming experiments, such as subcloning, PCR reactions, restriction digests, and transposon bombing, that were used to correct draft assemblies in the past.

We have typically employed SHIMS 3.0 on regions where we have prior knowledge of ampliconic structures that would render sequence assembly from short reads either very difficult (palindromes on primate X chromosomes) or impossible (the TSPY array on the human X chromosome). When sequencing a novel ampliconic target, it is often the case that the underlying genomic structures are known to be too complex for whole genome shotgun sequencing, but the exact repetitive structure is not known. In this case, we recommend sequencing a redundant tiling path of clones by SHIMS 2.0 first, and to deploy SHIMS 3.0 on the subset of clones that fail to assemble automatically from Illumina reads alone. This strategy provides the least expensive means for assembling the full range of ampliconic structures that we have encountered in vertebrate genomes, from tandem arrays of repeats tens of kilobases long to multi-megabase palindromes.

We adapted existing methods for generating ultra-long reads^17^ for use with pools of large-insert clones on the Oxford Nanopore Technologies (ONT) MinION platform. Successful generation of full-length reads requires intact DNA of high concentration and purity. We optimized our protocol to avoid unnecessary manipulations that could damage DNA or introduce contamination. We culture 24 clones separately, and then pool all cultures for DNA isolation, library preparation, and sequencing. In contrast to conventional plasmids, BACs and fosmids are present in only a single copy per host cell, and common reagents for increasing the efficiency of DNA precipitation, such as glycogen or SPRI beads, are incompatible with nanopore sequencing. We compensate for this by starting with large volume (∼15 ml) BAC and fosmid cultures to ensure that we harvest a sufficient amount of intact DNA. To preserve the integrity of high-molecular-weight (HMW) DNA, we handle it as little as possible, pipetting very slowly, using only wide-bore tips. We allow precipitated DNA to resuspend in water slowly over several days, rather than mixing by vortexing, or pipetting up-and-down. We have had the best results generating libraries from 7.5 to 15 µg of HMW DNA in 15 µl using the transposase-based RAD-004 library preparation kit from ONT. At these concentrations, solutions of HMW DNA will be extremely viscous, and it is difficult to measure the concentration precisely; some trial-and-error may be required to get the correct ratio of transposase to DNA, but we find that 0.5 µl of FRA for 15 µg is generally a good starting point to ensure that most BAC clones are cut only once. It is important to wait 45 minutes between loading the nanopore flow cell and starting the run, to allow time for full-length molecules to diffuse to the pores, otherwise the run will be dominated by shorter molecules. In contrast to our approach for generating Illumina reads, indexing or barcoding individual clones is not necessary, as full-length nanopore reads can be uniquely assigned to clones, even within the same amplicon.

Full-length nanopore reads transform clone finishing into a purely computational exercise. The tool chain for handling ultra-long nanopore reads is not yet fully mature, but it is developing rapidly. We rely on Minimap2 for alignments involving full-length reads^18^. This includes assigning nanopore reads to clones based on SHIMS 2.0 draft sequences, identifying full-length reads that start and end in vector sequence, and aligning a mix of long and short reads to generate a consensus. We use Racon for polishing the consensus sequence^19^, and SAMtools and custom scripts to manipulate read alignments^20^. We use Gap5 and Consed for visualizing discrepant bases and manually editing the consensus^21, 22^. While full-length reads guarantee the correct overall sequence structure, a variety of alignment artifacts may occur in clones with highly identical internal repeats. In this case, we find it is best to electronically split the clone sequence into individual repeat units, and correct each unit separately, before merging them together to create a finished clone sequence.

### Performance

The TSPY array on the Y chromosome is the largest and most homogeneous protein-coding tandem array in the human genome^16^, consisting of a 20.4-kb unit present in a highly variable number of copies, ranging from 11 to 72 per individual^23^. TSPY encodes a testis-specific protein, implicated in gonadoblastoma^24^, that regulates cell proliferation^25^; TSPY copy number is positively correlated with sperm count and sperm concentration^26^. As a demonstration of the expected performance of SHIMS 3.0, we fully resolved this array for the first time, using clones from the RP11 BAC library that we previously employed for our SHIMS 1.0 assembly of the male-specific region of the human Y chromosome^4^ (Fig. 2). This array spans 600 kb and contains 29 repeat units (Fig. 2a & b). We sequenced a redundant path of 19 clones and identified 94 sequence family variants that allowed us to select a non-redundant tiling path of 9 clones for finishing (Fig. 2c). On average, each unit differs from the others by 1 in 100 bases. The array encodes 14 distinct TSPY transcripts, encoding 10 different proteins, and it includes one pseudogene (Fig. 2d). We observed that at least 4 transcript variants were expressed in published testis RNA-seq datasets from other males^27^, indicating that multiple copies are expressed.

**Figure 2.**
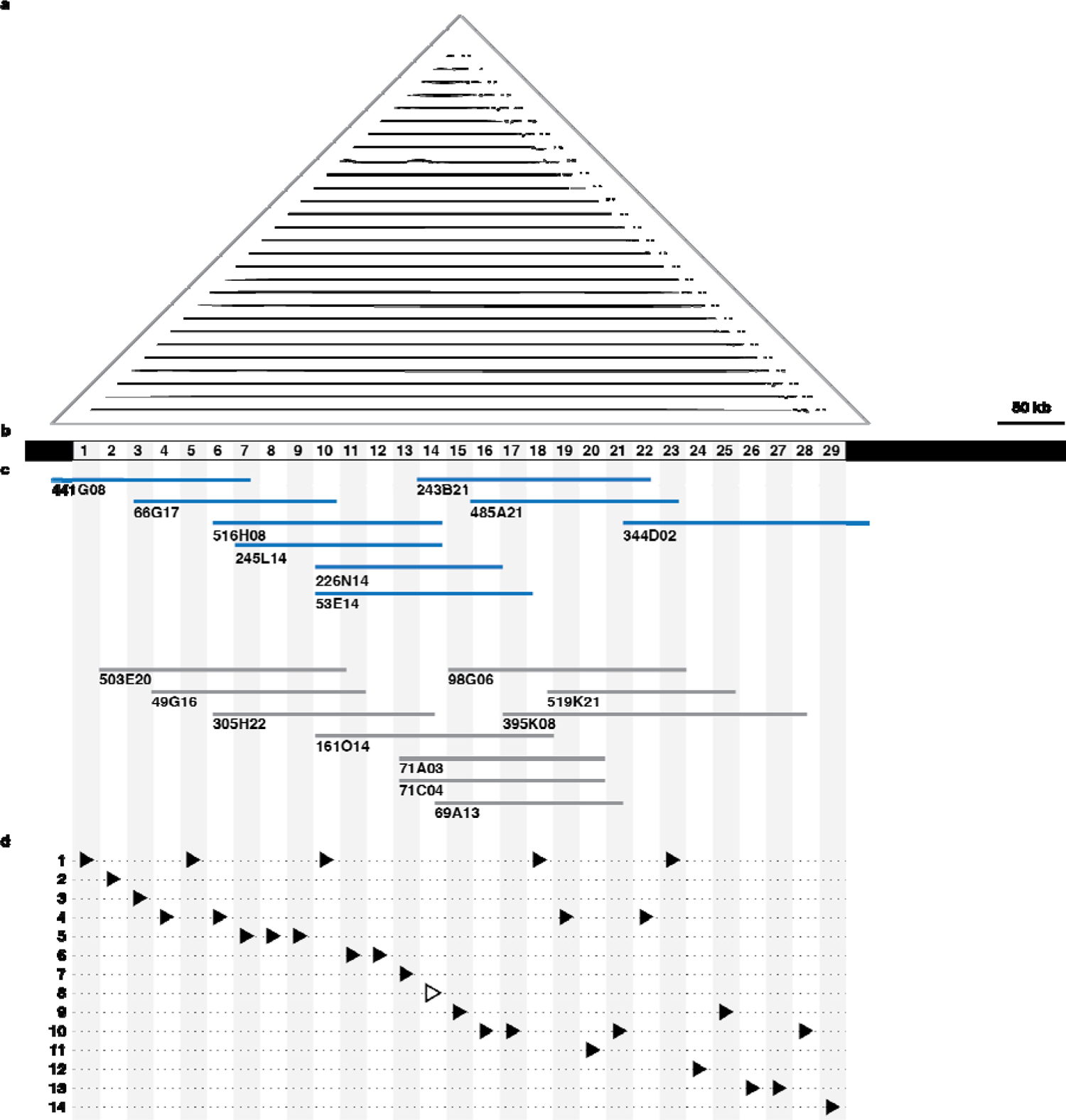
Structure of the Human TSPY array. a) Triangular dot plot of TSPY array region (AC279304) assembled using SHIMS3.0; each dot represents a 100 nucleotide perfect match between sequences within the array; scale bar: 50 kb. b) Schematic representation of the 29 repeat units of the TSPY array. c) Clones from the RP11 BAC library sequenced with SHIMS 2.0 and SHIMS 3.0 (blue) to obtain finished sequence, or SHIMS 2.0 alone (grey) to identify sequence family variants used to map the array. d) There are 14 distinct TSPY transcripts (triangles), including 1 pseudogene (open triangle).

Using only Sanger or Illumina reads, the presence of multiple amplicon copies within a single insert causes the clone assembly to collapse. The median BAC clone in the TSPY array contains 9 repeat units, making it impossible to accurately assemble even a single clone from this region using SHIMS 1.0 without months of manual finishing efforts at a genome center. By incorporating full-length nanopore reads, SHIMS 3.0 permits a small team – a technician and bioinformatics specialist – to finish these challenging sequencing targets. Our SHIMS 3.0 protocol can improve 24 SHIMS 2.0 draft sequences to finished quality in 2 weeks at a cost of $80 per clone. Despite the enormous reduction in staffing, cost, and time, the quality of finished sequence remains extremely high. We observe less than 1 error per megabase in overlaps between clones, on par with previous versions of SHIMS.

### Comparison with other methods

SHIMS produces de novo sequence assemblies with higher accuracy than any other technique, making it possible to produce accurate reference sequence of the most extreme repetitive regions, from ampliconic sequences like the nearly-perfect multi-megabase duplications on the mouse Y chromosome^9^, to the thousands of centromeric satellite repeats that form the centromere of the human Y chromosome^28^. This extremely high accuracy is due to the clone-based nature of SHIMS. Each clone represents a single long molecule that can be sequenced repeatedly, with comple mentary technologies, to generate an assembly that is accurate at the level of overall structure as well as the identity of individual nucleotides. This property makes it possible to identify and repeatedly verify the rare sequence family variants that distinguish ampliconic repeats, and build a high-confidence map from individual clones.

The impressive advances in single-molecule sequencing technologies that enabled SHIMS 3.0 have also increased the capabilities of whole genome shotgun approaches^29^. It is now routine to generate nanopore sequencing runs where half of the bases are in reads longer than 100 kb, so that interspersed repeats and smaller ampliconic structures can be spanned by a single long read. Whole genome shotgun with nanopore reads enabled the complete assembly of the human X chromosome from a single haplotype source, the CHM13hTERT cell line^30^. This effort required deep coverage from nanopore reads as well as from a broad array of complementary sequencing and mapping technologies, combined with manual review of structurally complex regions^30^. Error rates were still orders of magnitude higher than clone based strategies – 1 error in 10 kb in single-copy sequence, and 7 errors per kilobase in sequences present in more than one copy^30^. This elevated error rate in multi-copy sequence is due to the inherent difficulties of uniquely mapping short reads to long paralogous repeats; instead of reconstructing the true sequence, error correction with short reads blurs all paralogs together into an erroneous consensus. A second, orthogonal quality control measure indicates that shotgun sequencing with nanopore reads still lags behind clone-based approaches; 18% of CHM13 BAC sequences from segmental duplications and other difficult-to-assemble regions were missing from the whole genome shotgun assembly of CHM13hTERT^30^. While sequencing technologies continue to improve in read length and accuracy, clone-based approaches will continue to be relevant for generating highly accurate reference sequence, particularly in otherwise inaccessible ampliconic regions.

Relative to previous versions of SHIMS, SHIMS 3.0 greatly reduces the resources required to successfully generate finished sequence. Under SHIMS 1.0, the cost to produce draft sequence averaged about $5000 per clone, while finishing averaged around $4000. Weeks of bench experiments and many days of expert review were required to transform each low coverage (5-8x) Sanger draft sequence with frequent gaps into a complete and contiguous assembly. In SHIMS 2.0, we reduced the need for costly finishing activities by opting for much higher coverage (50-80x) in much cheaper Illumina reads. We encounter fewer coverage gaps at this higher depth, and also fewer library gaps because of differences in the library preparation protocol. We rely on sonication to provide random shearing, and amplify library fragments by PCR, as opposed to cloning fragments in E. coli. Although this deep and relatively even coverage ensured that wet-bench experiments were rarely required for finishing, structurally complex clones still required several days of expert review using an assembly editor like Consed. In the most complex cases, involving many paralogous repeats within a single clone, such as the TSPY array or centromeric satellite repeats, it was still impossible to completely resolve the correct structure.

By incorporating full-length nanopore reads from each clone, SHIMS 3.0 now makes it possible to assemble even the most internally repetitive clones. Full-length nanopore reads provide complete certainty about the overall clone structure; there is no doubt about the order and orientation of sequences, and no question about the copy number of complex repeats. This limits finishing activity to the simple matter of resolving the few remaining discrepancies between nanopore and Illumina reads. In most clones, this requires less than an hour of effort for even inexperienced finishers, and results in highly accurate sequences, with less than 1 error per megabase. Highly repetitive clones require more attention, but they can be resolved by a simple divide-and-conquer strategy, where each paralogous repeat is finished separately, with special attention to SFV sites, and then merged to create the full finished sequence. Correctly mapping short reads to repeated sequences becomes more difficult as the number of paralogs increases, increasing the chances that each paralogous repeat unit is blurred toward the consensus. In contrast to WGS strategies, in SHIMS 3.0, this blurring is confined to the boundaries of a single clone, and comparisons with neighboring clones can be used to resolve the position of paralogous SFVs. SHIMS 3.0 dramatically decreases the time, cost, and effort required to obtain finished sequence; using an optimized protocol for preparing HMW DNA in parallel from pools of BAC clones, a small team can finish 24 Illumina draft assemblies in 2 weeks for $80 per clone.

### Limitations of SHIMS 3.0

SHIMS 3.0 exceeds the capabilities of previous iterations of the SHIMS technique, providing access to the longest, most highly identical ampliconic sequences, as well as arrays of repeated sequences shorter than a single clone. However, SHIMS 3.0 shares two of the same limitations as previous versions of SHIMS and other clone-based approaches. First, the maximum size of BAC inserts limits SHIMS to resolving duplications with <99.999% identity. This limitation will remain until long-read technologies are able to surpass BAC sequencing in both read length and accuracy, or a reliable cloning technology emerges that exceeds the insert size of BACs. Second, SHIMS is limited to sequences that can be cloned into E. coli. Sequences that are toxic to E. coli are underrepresented in BAC and fosmid libraries. These library gaps can be resolved by directed efforts that avoid cloning in E. coli, like sequencing long-range PCR products^4^, or using the emerging selective sequencing (“ReadUntil”) capability of nanopore-based sequencers to enrich for reads flanking the gap.

Practitioners of SHIMS 3.0 also face new challenges due to their reliance on nanopore reads spanning 100-300 kb BAC inserts. Bioinformatics tools for aligning, visualizing, and editing reads of this length are not fully mature. SAM and BAM files both encode alignment details in the CIGAR format, however, the BAM format is limited to 65535 operations, which is frequently too few to encode the many transitions between matches, mismatches, insertions and deletions encountered in alignments of ultra-long nanopore reads^31^. Moreover, Consed does not reliably display alignments of reads longer than 1 kb. Our workaround has been to split SAM formatted alignments of nanopore reads into uniquely named sub-alignments every 1000 match operations, and convert the resulting SAM files to BAM format, which is accepted by Consed. This permits full visualization of nanopore reads alongside Illumina reads during finishing.

### Expertise

As with our previous protocol for SHIMS 2.0, we have designed the SHIMS 3.0 protocol to be carried out by a small team. A single technician can process 24 BAC clones from frozen stocks to nanopore sequencing libraries in 5 days with common molecular biology lab equipment. A bioinformatics specialist can set up a pipeline to identify full-length reads for each clone, generate a consensus sequence, automatically correct most errors using alignments with short reads, and manually review the resulting assembly for errors and identify SFVs. It is important to keep abreast of new developments in software for processing nanopore data, as all aspects from base-calling to alignment and error correction are continuously being improved.

## MATERIALS

### REAGENTS

- Ethyl Alcohol 200 Proof (Ethanol; Pharmco-Aaper, cat. no. 11000200) !CAUTION Ethyl Alcohol is flammable; keep away from flame when handling it.
- Tris Base (AmericanBio, cat. no. AB020000-05000)
- Hydrochloric Acid (HCl; VWR, cat. no. BDH3026-500MLP) !CAUTION Hydrochloric Acid is corrosive. Wear gloves and eye protection when handling it.
- Polyethylene glycol 8000(PEG-8000; Sigma-Aldrich, cat. no. P2139-2KG)
- Sodium Chloride (NaCl; AmericanBio, cat. no. AB01915-10000)
- Bacto-tryptone (BD Biosciences, cat. no 211705)
- Yeast Extract (BD Biosciences, cat. no 211929)
- Chloramphenicol (Sigma-Aldrich, cat. no. C0378-5G) !CAUTION Chloramphenicol powder is hazardous. Handle this reagent in ventilated fume hood with gloves and eye protection.
- Glycerol (EMD Millipore Corp., cat. no. 356350-1000ML)
- ZymoPURE II Plasmid Maxiprep Kit (Zymo Research, cat. no. D4203)
- Rapid Sequencing Kit (Oxford Nanopore Technologies, cat. no. SQK-RAD004)

### EQUIPMENT

- EZ-Vac Vacuum Manifold (Zymo Research, cat. no. S7000)
- MinION (Oxford Nanopore Technologies)
- NanoDrop 1000 Spectrophotometer (Thermo Fisher Scientific, cat. no. ND 1000)
- Centrifuge 5810 R (Eppendorf, cat. no. 00267023)
- Microcentrifuge 5425 (Eppendorf, cat. no. 5405000107)
- Vortex-Genie 2 (Scientific Industries, cat. no. SI-0236)
- Portable Pipet-Aid XP2 Pipette (Drummond, cat. no. 4-000-501)
- Eppendorf ThermoMixer (Eppendorf, cat. no. 5380000028)
- Beckman Coulter Avanti J-E centrifuge (Beckman Coulter, cat, no. A20698)
- Fisherbrand Low-Retention Microcentrifuge Tubes (Fisher Scientific, cat. no. 02-681-320)
- ART Barrier Specialty Pipette Tips, 1000, wide bore (Fisher Scientific, cat. no. 2069GPK)
- 50 ml Falcon Tube (Fisher Scientific, cat. no. 14-959-49A)
- Nalgene™ PPCO Centrifuge Bottles with Sealing Closure (Fisher Scientific, cat. no 3141-0250)
- Costar Assay Plate 96-well (Corning, cat. no. 3797)
- Nunc 96 DeepWell Plate (Thermo Fisher Scientific, cat. no. 278743)
- AirPore Tape Sheets (Qiagen, cat. no. 19571)
- Parafilm M film (Bemis, PM996)
- Aluminum Adhesive Foil (Bio-rad, cat. no. MSF1001)

### SOFTWARE

- minimap2 (https://github.com/lh3/minimap2)
- racon (https://github.com/lbcb-sci/racon)
- samtools (http://www.htslib.org)
- tg_index (http://staden.sourceforge.net)
- gap5 (http://staden.sourceforge.net)
- (optional) consed (http://www.phrap.org/consed/consed.html)

### REAGENT SETUP

**70% (vol/vol) Ethanol**

Mix 30 ml of 100% (vol/vol) ethanol with 70 ml of of distilled, deionized water (ddH_2_O).

▴CRITICAL 70% (vol/vol) ethanol should be prepared on the day of the experiment.

**1M Tris-Cl, pH 8.5**

Dissolve 121 g of Tris base in 800 ml of ddH_2_O. Adjust pH to 8.5 with concentrated HCl, then adjust volume with ddH_2_O to 1 l. 1M Tris-Cl can be prepared in advance and stored at room temperature (22 °C) for up to a year.

**10 mM Tris-Cl, pH 8.5**

Mix 0.5 ml of 1M Tris-Cl with 49.5 ml of ddH_2_O. This solution can be prepared in advance and stored at room temperature for up to a year.

**PEG buffer (18% PEG(wt/vol)/1M NaCl Solution)**

Add 135 g of PEG-8000 powder into 1 l bottle. Add 150 ml of 5M NaCl, 7.5 ml of Tris-HCl, 1.5 ml of 0.5M EDTA and 450 ml of ddH_2_O to make PEG buffer. Store at room temperature for up to 1 year.

**2X LB**

Add 20 g of bacto-tryptone, 10 g of yeast extract, and 20 g of NaCl to ddH_2_O, and adjust the volume to 1 l. Mix well with a magnetic stirrer. After mixing, distribute 500-ml aliquots into 1 l bottles. Cap loosely, prewarm to 50 °C, and autoclave for 20 min on liquid cycle. Store at room temperature for up to 1 year.

**Chloramphenicol**

Dissolve 0.34 g of chloramphenicol into 10 ml of 100% (vol/vol) ethanol. Chloramphenicol stock can be stored at −20 °C for up to 1 year.

**80% (vol/vol) Glycerol solution**

Add 400 ml of glycerol to a graduated cylinder; adjust the volume to 500 ml with ddH_2_O. Seal the cylinder with Parafilm M film, and mix by inversion. Transfer to a bottle, and autoclave for 20 min in liquid cycle. This solution can be prepared in advance and stored at room temperature for up to 1 year

### PROCEDURE

#### Pick Clones and Grow Cultures ⍰TIMING 18 h

1| Fill each well of a Nunc 96 DeepWell plate with 1.9 ml of 2X LB containing 34µg/ml chloramphenicol.

▴CRITICAL STEP Rich media (2X LB) is appropriate for single-copy plasmids like BACs or fosmids, which use chloramphenicol resistance as a selectable marker.

2| Use a clean pipette tip to scrape the surface of a frozen glycerol stock and drop the tip directly into the DeepWell plate to inoculate a well. Inoculate each sample 8 times for a total of 15.2 ml/sample. 24 samples in total for each library prep.

▴CRITICAL STEP Start with a glycerol stock of a clone verified to be correct by PCR for a known sequence-tagged site or previous sequencing experiments (e.g. cultures created during Step 7 of the SHIMS 2.0 protocol).

3| Seal plates with AirPore Tape Sheets and place at 37 °C for 16-17 h, shaking at 220 RPM.

▲CRITICAL STEP Overgrowth of cultures (cell density > 3 x 10^9^cells per ml) will decrease yield of BAC DNA.

Glycerol Stock Plate ⍰TIMING 30 min

4| Dispense 150 μl of 80% (vol/vol) glycerol solution into two rows of a Costar Assay Plate.

5| Transfer 150 μl of each sample culture from Step 3 to a corresponding well of the assay plate and mix by pipetting up and down 20 times.

6| Seal the glycerol stock plate with aluminum adhesive foil. 7| Store the glycerol stock plate at −80 °C.

Pooling Clones ⍰TIMING 1-2h

8| Pour overnight cultures from Step 3 into a large beaker to combine pool.

9| Divide pooled culture into two 250 ml Nalgene bottles and spin down culture at 6000g for 30 min at 4 °C.

10| Remove media by pouring into a waste-collecting container. Be careful not to disturb the pellets.

LJ PAUSE POINT Store at −20 °C for up to a week.

#### Alkaline Lysis ⍰TIMING 1-2 h

11| Add 7 ml of ZymoPURE P1 (Red) to each bacterial cell pellet and resuspend completely by pipetting. Combine into one bottle when cells are completely resuspended.

12| Add 14 ml of ZymoPURE P2 (Green) and immediately mix by gently inverting the tube 6 times.

▴CRITICAL STEP Do not vortex! Let sit at room temperature for 3 min. Cells are completely lysed when the solution appears clear, purple, and viscous.

13| Add 14 ml of ZymoPURE P3 (Yellow) and mix gently but thoroughly by inversion.

▴CRITICAL STEP Do not vortex! The sample will turn yellow when the neutralization is complete, and a yellowish precipitate will form.

14| Ensure the plug is attached to the Luer Lock at the bottom of the ZymoPURE Syringe Filter. Place the syringe filter upright in a tube rack and load the lysate into the ZymoPURE Syringe Filter and wait 8 min for the precipitate to float to the top.

15| Remove the Luer Lock plug from the bottom of the syringe and place it into a clean 50 ml conical tube. Place the plunger in the syringe and push the solution through the ZymoPURE Syringe Filter in one continuous motion until approximately 33-35 ml of cleared lysate is recovered. Save the cleared lysate!

16| Add 14ml ZymoPURE Binding Buffer to the cleared lysate from step 5 and mix thoroughly by inverting the capped tube 10 times.

17| Ensure the connections of the Zymo-Spin V-P Column Assembly are finger-tight and place onto a vacuum manifold.

18| With the vacuum off, add the entire mixture from step 6 into the Zymo-Spin V-P Column Assembly, and then turn on the vacuum until all the liquid has passed completely through the column.

19| Remove and discard the 50 ml reservoir from the top of the Zymo-Spin V-P Column Assembly.

20| With the vacuum off, add 5 ml of ZymoPURE Wash 1 to the 15 ml Conical Reservoir in the Zymo-Spin V-P Column Assembly. Turn on the vacuum until all the liquid has passed completely through the column.

21| With the vacuum off, add 5 ml of ZymoPURE Wash 2 to the 15ml Conical Reservoir. Turn on the vacuum until all the liquid has passed completely through the column. Repeat this wash step.

22| Remove and discard the 15 ml Conical Reservoir and place the Zymo-Spin V-P Column in a Collection Tube. Centrifuge at ≥10,000g for 1 minute, in a microcentrifuge, to remove any residual wash buffer.

23| Pre-warm 450 μl of 10 mM Tris-Cl to 50 °C. Transfer the column into a clean 1.5ml microcentrifuge tube and add the 450 μl of 10 mM Tris-Cl directly to the column matrix. Wait 10 min, and then centrifuge at ≥ 10,000g for 1 minute in a microcentrifuge.

24| Add 450 μl of PEG buffer to the tube containing sample. Mix by flicking and rotating the 1.5 ml microcentrifuge tube.

25| Centrifuge at ≥10,000g for 30 min at 4 °C, in a microcentrifuge. 26| Remove supernatant from the tube without disturbing the pellet. 27| Add 1 ml of 70% EtOH and spin for 10 min at 4 °C.

28| Repeat step 27 and 28.

29| Remove supernatant and any left over 70% EtOH from microcentrifuge tube. 30| Air dry for 10 min or until no visible liquid is left in the tube.

▴CRITICAL STEP Do not overdry the pellet. 31| Dissolve DNA pellet in 18 µl 10 mM Tris-Cl.

32| Store DNA at 4 °C for several days until pellet completely dissolves into solution.

33| Check DNA concentration and quality with Qubit or NanoDrop. Aim for concentration ≥ 1 μg/μl, an A260/280 ratio ∼ 1.8, and an A260/230 ratio between 2.0 and 2.2.

?TROUBLESHOOTING

### MinION Library prep lilTIMING 30 Minutes

34| Adjust sample concentration from step 33 to 1 µg/µl with 10 mM Tris-Cl.

?TROUBLESHOOTING

35| Using a wide bore pipette tip, slowly aspirate 15 µl into a low retention microcentrifuge tube.

▴CRITICAL STEP Pipetting can shear fragile high-molecular-weight DNA. Pipette slowly using wide bore pipette tips.

36| In a separate microcentrifuge tube, add 0.5 µl FRA to 4.5 µl 10 mM Tris-Cl. Flick the tube to mix well.

▴CRITICAL STEP The FRA solution is included in the Rapid Sequencing Kit.

37| Add the diluted FRA solution from step 36 into sample tube from step 35. 38| Gently flick the tubes a few times to mix.

▴CRITICAL STEP Vortexing can shear fragile high-molecular-weight DNA. Flick solutions to mix.

39| Incubate sample on 30 °C heat block for 35 seconds, then move the tube to 80

°C heat block. Incubate for 2 min at 80 °C.

40| Remove the tube from heat block and incubate on ice for 1 minute, then move the tube off the ice. Equilibrate to room temperature, about 1 minute.

41| While the sample is equilibrating to room temperature, add 4.5 µl 10 mM Tris-Cl to 0.5 µl of RAP. Flick to mix well.

▴CRITICAL STEP The RAP solution is included in the Rapid Sequencing Kit. 42| Add RAP dilution from step 41 into sample tube. Slowly flick the tube a few times to mix. Keep the sample at room temperature before loading.

▴CRITICAL STEP Vortexing can shear fragile high-molecular-weight DNA. Flick solutions to mix.

MinION Library loading ⍰TIMING 30 Minutes

43| Add 30 µl of FLT to one tube of FLB, to make flush buffer, according to the Rapid Sequencing Kit instructions. Vortex the solution to mix, then centrifuge briefly.

▴CRITICAL STEP The FLT and FLB solutions are included in the Rapid Sequencing Kit.

44| Perform QC on a new MinION flow cell to check available pores and ensure that a sufficient number of pores are present. If there are fewer than 800 available pores, place the flow cell in storage and use a new MinION flow cell.

45| Use a P1000 pipette to remove about 20-30 µl of storage buffer from priming pore. Load 800 µl flush buffer via the pore slowly. Wait 5 min.

46| Lift SpotON cover and load 200 µl flush buffer slowly. Try to dispense at a speed where each bead of liquid is siphoned into the SpotON port as soon as it is visible.

47| Add 34 µl SQB and 15 µl ddH_2_O to the sample tube from step 42.

▴CRITICAL STEP The SQB solution is included in the Rapid Sequencing Kit. 48| Flick the tube gently to mix, then centrifuge briefly to collect library to the bottom of the tube.

▴CRITICAL STEP Vortexing can shear fragile high-molecular-weight DNA. Flick solutions to mix.

49| Slowly aspirate 75 µl of library with a wide bore tip. Very slowly, load the library into SpotON pore drop by drop.

▴CRITICAL STEP Pipetting can shear fragile high-molecular-weight DNA. Pipette slowly using wide bore pipette tips.

50| Close both priming pores and put the SpotON cover back onto the pore. 51| After loading the library, leave the flow cell on bench for 45 min before starting the run.

Wait at least 45 minutes between loading the flow cell and starting the run. This allows time for full-length molecules to diffuse to the pores. Starting the run earlier will favor the sequencing of shorter molecules.

?TROUBLESHOOTING

### Demultiplex Reads ⍰TIMING 30 Minutes

`draft_clones.fa`

52| Prepare file of draft clone sequences in fasta format:

▴CRITICAL STEP When concatenating draft sequence assemblies, ensure that each sequence has a unique name.

53| Prepare file of vector sequence in fasta format: `vector.fa`

54| Download fastq formatted reads from the device running MinION control software: `nanopore.fq`

55| Align nanopore reads to file of draft sequences to assign nanopore reads to clones by best match:

?TROUBLESHOOTING

`minimap2 -x map-ont draft_clones.fa nanopore.fq | sort`

`-r -n -k 10 | awk ‘!seen[$1]++’ > best_clone_match.paf`

`grep clone_name best_clone_match.paf | cut -f 1`

`>clone_name.txt`

`grep -A 3 -f clone_name.txt nanopore.fq | grep -v “^-- $” > clone_name.nanopore.fq`

Identify Full-length Reads ⍰TIMING 30 Minutes

▴CRITICAL We have automated Steps 56-65 with a custom Perl script (available at https://github.com/dwbellott/shims3_assembly_pipeline/), but the workflow is described below to allow for direct use of the individual software tools or substitution of alternative tools.

56| For each clone, align nanopore reads to file of vector sequence:

`minimap2 -x map-ont vector.fa clone_name.nanopore.fq –o clone_name.vector.paf`

57| Search for reads that begin and end with high-quality matches to vector sequence on the same strand – these are full-length reads.

`cut -f 1,5,6 clone_name.vector.paf | sort | uniq -c | sed ‘s/^//’ | grep “^2” | cut -f 2 -d ‘>clone_name.2x.txt’`

`awk ‘$2 - $3 < $7 && $12 == 60’ clone_name.vector.paf | cut -f 1,5,6 | grep -f clone_name.2x.txt`

`>clone_name.2x.right.txt`

`awk ‘$4 < $7 && $12 == 60’ clone_name.vector.paf | cut`

`-f 1,5,6 | grep -f clone_name.2x.right.txt | cut -f 1 | sort | uniq >clone_name.fl.txt`

58| For each clone, generate a fastq file of full-length reads, as well as a fasta file of the longest full-length read to use as a scaffold for final assembly.

?TROUBLESHOOTING

`grep -A 3 -f fl.txt clone_name.nanopore.fq | grep -v “^--$” > clone_name.fl.fq`

grep -A 1 ‘head -n 1 clone_name.fl.txt’ nanopore.fq | sed ‘s/^\@.*/\>clone_name/’ >clone_name.longest.fl.fa

Generate Consensus Sequence ⍰TIMING 30 Minutes

59| Polish the longest read twice, using the other full-length nanopore reads:

`minimap2 -x map-ont clone_name.longest.fl.fa clone_name.fl.fq >clone_name.longest.fl.paf`

`racon clone_name.fl.fq clone_name.longest.fl.paf clone_name.longest.fl.fa`

`>clone_name.longest.fl.racon.fa`

`minimap2 -x map-ont clone_name.longest.fl.racon.1.fa clone_name.fl.fq >clone_name.longest.fl.racon.paf`

`racon clone_name.fl.fq clone_name.longest.fl.racon.1.paf clone_name.longest.fl.racon.1.fa`

`>clone_name.fl.consensus.fa`

60| Gather up Illumina, nanopore, and (if available) PacBio reads for each clone.

`cat clone_name.illumina.forward.fq`
`clone_name.illumina.reverse.fq`
`clone_name.illumina.single.fq`
`clone_name.nanopore.fq`
`clone_name.pacbio.fq >>`
`clone_name.allreads.fq`

61| Polish the nanopore consensus sequence, using both long and short reads.

`minimap2 -x asm20 clone_name.nanopore.consensus.fa`
`clone_name.illumina.single.fq >>`
`clone_name.polish.1.paf`

`minimap2 -x sr clone_name.nanopore.consensus.fa`
`clone_name.illumina.forward.fq`

`clone_name.illumina.reverse.fq >>`
`clone_name.polish.1.paf`

`minimap2 -x map-ont clone_name.nanopore.consensus.fa`
`clone_name.nanopore.fq >>`
`clone_name.polish.1.paf`

`minimap2 -x map-pb clone_name.nanopore.consensus.fa`
`clone_name.pacbio.fq >>`
`clone_name.polish.1.paf`

`racon clone_name.allreads.fq`
`clone_name.polish.1.paf`
`clone_name.nanopore.consensus.fa`

`>clone_name.polish.1.fa`

62| Repeat Step 61 four more times, for a total of 5 rounds of polishing

63| Align reads one last time to generate SAM format alignments suitable for assembly editors.

#### The BAM file format cannot accommodate CIGAR strings with greater than 65535 operations. Alignments involving nanopore reads spanning the full length of a BAC clone will exceed this limit. We strongly recommend storing alignments in SAM or CRAM format to avoid the loss of detailed alignment information

minimap2 -x asm20 -a -L --sam-hit-only -R ‘@RG\tID:S\tSM:S\tPL:ILLUMINA’ clone_name.polish.5.fa clone_name.illumina.single.fq | samtools sort -O SAM -

`>clone_name.single.sorted.sam`

minimap2 -x sr -a -L --sam-hit-only -R ‘@RG\tID:FR\tSM:FR\tPL:ILLUMINA’ clone_name.polish.5.fa

`clone_name.illumina.forward.fq clone_name.illumina.reverse.fq | samtools sort -O SAM -`

`>clone_name.paired.sorted.sam`

minimap2 -x map-pb -a -L --sam-hit-only -R ‘@RG\tID:P\tSM:P\tPL:PACBIO’ clone_name.polish.5.fa clone_name.pacbio.fq | samtools sort -O SAM -

`>clone_name.pacbio.sorted.sam`

minimap2 -x map-ont -a -L --sam-hit-only -R ‘@RG\tID:N\tSM:N\tPL:PACBIO’ clone_name.polish.5.fa clone_name.nanopore.fq | samtools sort -O SAM -

`>clone_name.nanopore.sorted.sam`

64| Combine alignments

`samtools merge -f clone_name.allreads.sorted.sam clone_name.single.sorted.sam clone_name.paired.sorted.sam clone_name.pacbio.sorted.sam clone_name.nanopore.sorted.sam`

65| Generate database for Gap5

We now recommend Gap5 over Consed, because Gap5 natively supports loading data directly from SAM files and displaying full-length nanopore reads. It is possible to split SAM alignments of full-length nanopore reads into smaller fragments that can be encoded in a BAM file and displayed by Consed without loss of information. For those who wish to use

Consed, we implement this work-around in a custom Perl script (available at https://github.com/dwbellott/shims3_assembly_pipeline/).

`tg_index -o clone_name.g5d -p −9 -s`

`clone_name.allreads.sorted.sam`

Finishing ⍰TIMING 0-8 h

66| Open the assembly in Gap5:

`gap5 clone_name.g5d`

67| Select ‘Edit Contig’ from the ‘Edit’ menu.

68| Resolve discrepancies between Illumina reads and full-length nanopore reads (Fig. 3).

**Figure 3.**
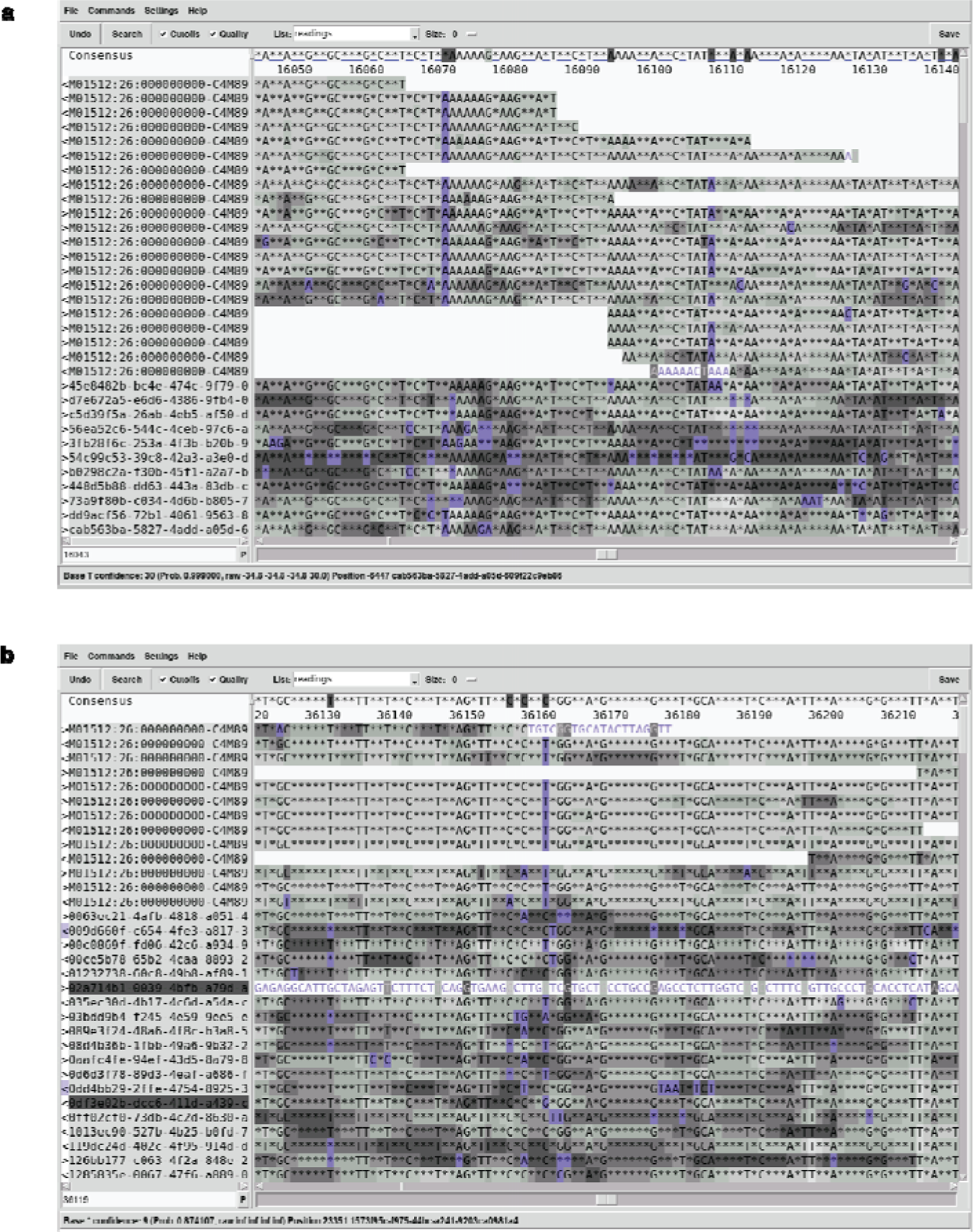
Editing clone assemblies in Gap5. Screenshots from Gap5 with reads sorted by technology (Illumina on top; nanopore on bottom), showing two instances where errors in the consensus can be resolved by correcting to the consensus of the Illumina reads: a) frequent insertion and deletion errors at homopolymer runs, and b) more rare substitution errors.

In Gap5, it is not possible to directly edit the consensus sequence. Instead, indicate which readings are authoritative by marking bases as high quality with the ‘]’ key, and the consensus will automatically update.

We usually resolve the consensus in favor of the Illumina reads. The vast majority of discrepancies between these technologies occur at homopolymer repeats, where nanopore reads are especially prone to insertion and deletion errors (Fig. 3a). More rarely, we encounter systematic errors in nanopore base calling that generate a consensus that is not supported by any Illumina read.

▴CRITICAL STEP We resolve disagreements among Illumina reads in favor of the consensus of full-length nanopore reads. In clones that contain duplicated sequences, short Illumina reads can be mapped to the wrong repeat unit, but full-length nanopore reads are not subject to this artifact, and will usually have the correct base at each SFV.

69| Resolve SSRs by realigning reads around the SSR region. Select reads by clicking on their names on the left hand side of the edit window, and choose ‘Realign Selection’ from the ‘Command’ menu.

▴CRITICAL STEP Stutter noise from replication slippage in SSRs causes divergent reads and low-quality base calls. In some cases, unambiguous resolution of these repeats may not be possible, and they should be annotated as unresolved in Step 71.

70| Remove any vector-sequence contamination at the ends of the clone. In the Gap5 edit window, use the ‘Search’ button to search the consensus sequence for the sequences at the cloning site of your vector. Trim away the vector sequence outside of the restriction sites used to generate your clone library (usually EcoRI, BamHI, or MboI).

71| Annotate any remaining ambiguities in the clone sequence (e.g., unresolved simple sequence repeats, where neither Illumina or nanopore reads are completely accurate) by compiling a feature table^32^, which will be useful when finished clone sequences are submitted to GenBank.

?TROUBLESHOOTING

⍰TIMING

Steps 1-3, pick clones and grow cultures: 18 h

Steps 4-7, glycerol stock plate: 30 min

Steps 8-10, pooling clones: 1-2 h

Steps 11-33, alkaline lysis: 1-2 h

Steps 34-42, MinION library prep: 30 min

Steps 43-51, MinION library loading: 30 min

Steps 52-55, demultiplex reads: 30 min

Steps 56-58, Identify full-length reads: 30 min

Steps 59-65, generate consensus sequence: 30 min

Steps 66-71, finishing: 0-8 h

?TROUBLESHOOTING

#### Troubleshooting advice can be found in Table 1

### Anticipated Results

We typically pool 24 clones for a single MinION run, generating about 300,000 reads with a read n50 of 20 kb, and a total of about 1.5 Gb of sequence data. Each clone typically receives 1-5% of the total reads. Occasionally some clones will have no reads; this usually indicates that the culture of the clone (Steps 1-3) has failed (see troubleshooting information for Step 55).

**Table 1.** Troubleshooting Table.

| Step | Problem | Possible Reason | Solution |
| --- | --- | --- | --- |
| 33 | Low DNA concentration | Culture undergrowth or overgrowth | Check culture OD600 is between 0.2-0.35 |
|  |  | Incomplete Lysis | Make sure to thoroughly mix the solution until the color is uniform |
|  |  | Incomplete neutralization | Solution from step 13 should not appear viscous and precipitate should float to the surface |
|  |  | Incomplete DNA elution | Pre-warm elution buffer to 50°C |
| 34 | Concentration varies when checking with NanoDrop or Qubit | DNA is not completely mixed | After adjusting concentration from step 33, leave DNA solution on a heated shaker at the gentlest setting at 50°C until DNA is completely mixed |
| 51 | Pores decrease rapidly | Impure DNA sample | Re-check DNA concentration. Extract DNA again if NanoDrop and Qubit results are discordant, $260/280 < 1.7$, $260/280 > 2.0$, $260/230 < 2.0$, or $260/230 > 2.2$ |
|  |  | Bubbles introduced during loading | Pipette very slowly and take care not to introduce bubbles during flow cell priming and library loading |
| 55 | No reads for one or more clones | Clone culture failed | Regrow and add to the next run, or replace the clone with another |
|  |  |  | Regrow the clone for an additional round of sequencing |
|  |  | Bookkeeping error; some common bookkeeping errors result from transposing digits, rotating a plate by 180°, or contamination from a clone in an adjacent well | Resolve bookkeeping error, and rerun a new clone or replace with another clone |
| 58 | Low fraction of long reads | FRA treatment time too long | Promptly heat-inactivate FRA at 35 seconds |
|  |  |  | Adjust the FRA incubation time below 35 seconds |
|  |  | Shearing during library prep | Use wide-bore tips for all mixing and loading steps |
| 71 | Clone sequence is shorter than expected or missing known sequence | Deletion during culture | Regrow the clone from the original culture or another library copy, and replace with the alternate clone |
|  |  | Sequence toxic to *E. coli* | Close the gap by long-range PCR or region-specific extraction |

Expect to obtain 3-10 full-length reads per clone. Because of the high rate of insertions and deletions in individual nanopore reads, full-length reads may differ in length by 10 kb or more. Occasionally, a clone will have no reads that start and end in vector sequence, but the clone length will be apparent from a peak in the tail of the distribution of read lengths. It may still be possible to reconstruct a full-length consensus sequence by rotating one of these putative full-length reads to place the vector sequence at the beginning. However, we do not recommend this procedure for internally repetitive clones, particularly tandem arrays. Instead, sequence the clone again, and use these ambiguous reads to help polish the consensus.

## Acknowledgements

This work was supported by the Howard Hughes Medical Institute, and generous gifts from Brit and Alexander d’Arbeloff and Arthur W. and Carol Tobin Brill

## Author Contributions

D.W.B., H.S., J.F.H., and D.C.P. designed the study. D.W.B., T.-J.C., and E.K.J. developed the experimental methods. D.W.B. wrote the scripts for computational analysis. T.J.C. carried out the sequencing of the TSPY region. H.S. assembled the TSPY region. D.W.B., T.-J.C., and D.C.P. wrote the manuscript.

## Data Availability

Data that support the findings of this study have been deposited in Genbank with the accession codes AC279301, AC279302, AC279304, AC279305, AC279306, AC279307, AC279309, AC279311, AC279312, and AC279315.

## Code Availability

We have automated Steps 56-65 with a custom Perl script, available at: https://github.com/dwbellott/shims3_assembly_pipeline/

